# RNA-guided contrastive learning enhances patient-level prediction from histology

**DOI:** 10.64898/2026.09.24.754149

**Authors:** Anqi Zou, Elain Huang, Qiyou Wu, Mollie E. Barnard, Chao Zhang

## Abstract

Predicting molecular receptor status, including estrogen receptor (ER), progesterone receptor (PR), and human epidermal growth factor receptor 2 (HER2), directly from routine hematoxylin and eosin (H&E)-stained whole-slide images (WSIs) could reduce reliance on costly immunohistochemistry, but image-only models have no access to the transcriptomic programs that define these subtypes. Here we show that coupling a whole-slide vision transformer (GigaPath) with bulk RNA-sequencing profiles through a training-time-only contrastive objective improves receptor-status classification from WSIs alone, without requiring any RNA-seq data at inference. Rather than training a dedicated bulk-transcriptomics encoder, we purpose the gene-sentence encoding and pretrained text encoder from BioBERT^1^ and apply them to bulk RNA-seq profiles from The Cancer Genome Atlas breast cancer cohort (TCGA-BRCA). Under five-fold cross-validation, this RNA-guided pre-training consistently improved AUROC for ER, PR, and HER2 relative to a contrastive-pre-training-free baseline and reproducibly introduced receptor-relevant structure into the learned slide representation. However, evaluation on independent external cohorts revealed a striking, receptor-dependent divergence: gains for ER and PR, both governed by broad, multi-gene luminal transcriptional programs, generalized robustly, whereas the apparent AUROC improvement for HER2, driven by focal ERBB2 amplification rather than a coordinated transcriptional signature, masked a severe loss of classification sensitivity at the operating threshold. These findings establish training-time transcriptomic supervision as a practical route to enhancing histology-only molecular subtyping, while showing that gene-selection strategy, and not model architecture alone, determines which molecular alterations such supervision can teach a vision model to recognize.

## Introduction

Breast cancer has surpassed lung cancer as the most commonly diagnosed cancer globally^2^. It remains one of the most prevalent diseases due to its heterogeneous nature and resistance to therapy^3^. In clinical practice, the diagnostic process begins with pathologists evaluating hematoxylin and eosin (H&E)-stained whole-slide images (WSIs) to determine the type, grade, and extent of the breast specimen^4^. This is followed by biomarker testing using immunohistochemistry (IHC) to determine the expression status of the estrogen receptor (ER), progesterone receptor (PR), and human epidermal growth factor receptor 2 (HER2)^5,6^. These clinical markers are further categorized into intrinsic subtypes such as Luminal A, Luminal B, HER2-enriched, and basal-like, defined by genomic assays such as the Prediction Analysis of Microarray 50 (PAM50)^7^. Integrated histopathological and molecular profiling guides personalized treatment, ranging from endocrine and targeted therapies to immunotherapy and emerging adjunctive modalities like photodynamic therapy^8^. While this diagnostic workflow remains the clinical benchmark, it is limited by its demand for specialized expertise in the pathology workforce, which is strained by an ageing demographic, increased workloads, and increased complexity. Enabling accurate subtype classification directly from WSIs would expand the volume of cases available for subtype-specific research. Therefore, it is of great interest to develop a computational pipeline to deliver rapid, standardized, and reproducible subtype classification.

Despite advances in computational pathology, WSIs present a significant challenge due to their gigapixel scale, which exceeds the capacity of standard machine learning architectures. Consequently, many existing models rely on patch-level analysis, lacking high-level features such as spatial relationships between cells, tumor staging, and complex microscopic patterns. Furthermore, early models relied on imaging data, which lacks the underlying genetic profiles essential for molecular subtyping. The integration of histopathological morphology and transcriptomic expression is motivated by the relationship between a tumor’s genetic profile and its physical tissue morphology. While WSIs provide information on tissue structure, cellular morphology, and the immune microenvironment, they do not reveal the patient’s genomic landscape. Conversely, transcriptomic profiles provide information on the transcriptome, capturing gene expression patterns that define molecular and intrinsic subtypes. By using both modalities, a machine learning model can learn to identify morphological signatures that correlate with specific transcriptomic states, bridging the gap between imaging and ‘omics’ data.

Recent developments, such as OmiCLIP, a visual-omics foundation model^9^, have demonstrated that integrating H&E images with spatial transcriptomics (ST) using tissue patches from Visium data significantly enhances model performance, providing important evidence that cross-modal contrastive learning between histopathology and transcriptomics can yield biologically informed image representations. However, because Visium-based models are structured to process tissue as discrete patches, they do not align with the clinical goal of evaluating the WSI as a whole. Moreover, ST remains a prohibitively expensive assay. The cost of processing a single slide can exceed $2,500, creating a significant barrier to the large-scale data acquisition required for robust model training and contradicting the objective of expanding the volume of cases available for subtype-specific research. In contrast, bulk RNA-sequencing (RNA-seq) provides a more viable and scalable alternative: it is cost-effective, widely available in extensive public databases such as TCGA, and uniquely suited for a slide-level approach that treats the WSI as a singular, comprehensive diagnostic unit. Rather than developing a dedicated bulk RNA-seq encoder for this purpose, we note that existing transcriptomic encoding strategy, representing a transcriptomic profile as a ranked “gene sentence” of its most highly expressed genes, processed by a pretrained text encoder, does not depend on the spatial, spot-level structure of Visium data, and can be applied directly to bulk RNA-seq profiles by treating each patient’s bulk transcriptome as a single sample analogous to one Visium spot. This motivates our use of gene-sentence encoding scheme and pretrained text encoder as the transcriptomic branch of our framework, allowing us to couple bulk RNA-seq-derived representations with WSI-derived morphological features for slide-level, cost-effective subtype classification, without requiring a dedicated bulk RNA-seq foundation model.

In this work, we present an end-to-end computational pipeline that integrates histopathological WSIs with RNA-seq data for breast cancer subtype classification. By leveraging a contrastive learning framework, our approach overcomes the gigapixel scale of WSIs to provide a slide-level analysis that aligns with clinical diagnostic standards. We validated our model using public and private datasets, demonstrating that the multi-modal analysis approach of morphological features and transcriptomic profiles outperforms single-modality methods. This pipeline offers a cost-effective and reproducible alternative for identifying molecular subtypes, expanding the accessibility of subtype-specific research.

## Results

### Method overview

We present a training-time multi-modal contrastive learning framework that couples whole-slide H&E histopathology images with bulk RNA-seq profiles to enhance molecular subtype prediction in breast cancer, while requiring only an H&E-stained WSI at inference. Unlike the original purpose of existing gene-sentence encoding scheme, which was designed for spot-level spatial transcriptomics, we redefined the architecture to adapt it for WSI. By aligning a pretrained slide-level vision encoder^10^ with a transcriptomic (text) encoder via contrastive learning on patient-matched WSI-RNA pairs, our approach injects transcriptomic-informed molecular context into the visual representation without requiring RNA-seq data at test time, preserving a strictly unimodal inference pipeline. To validate whether this RNA-guided contrastive pre-training improves downstream task performance, we implement classification of ER, PR, and HER2 status, comparing against a vanilla, contrastive-pre-training-free GigaPath baseline evaluated under an identical protocol. We also examine the resulting models beyond aggregate AUROC, including the structure of the learned embedding space and the sensitivity-specificity trade-offs of the resulting classifiers, to characterize both the benefits and the limitations of this training-time-only molecular supervision. Together, these results indicate that bulk transcriptomic profiles are an effective source of training-time-only supervision for WSI-based molecular subtyping, using only publicly available data and without requiring a dedicated bulk RNA-seq foundation model.

### Data collection and pre-processing

All training data were obtained from The Cancer Genome Atlas (TCGA), specifically the breast cancer cohort (TCGA-BRCA). The dataset includes 1,098 samples, each comprising H&E-stained WSIs, bulk RNA-seq data, and corresponding molecular subtype labels (**Table 1**). Molecular subtypes were obtained from previous studies^11,12^, where tumor samples from the same cases were analyzed using immunohistochemistry (IHC) to determine receptor status. Specifically, samples were stained for ER, PR, and HER2. These biomarkers are commonly used in breast cancer subtype classification and clinical decision-making. In addition to receptor status, tumor samples were categorized into intrinsic molecular subtypes using the PAM50 gene expression classifier, which classifies tumors based on gene expression patterns rather than protein markers. The PAM50 classifier defines five major breast subtypes: Luminal A, Luminal B, HER2-enriched, Basal-like, and Normal-like. In total, the dataset includes 1,755 samples with ER labels, 1,531 samples with PR labels, and 377 samples with HER2 labels. Among the 844 samples with PAM50 subtype annotation, 380 samples are classified as Luminal A, 208 as Luminal B, 98 as HER2-enriched, 142 as Basal-like, and 15 as Normal-like. To evaluate model performance and ensure robustness, both the IHC receptor labels and PAM50 subtype labels were partitioned into five folds for 5-fold cross-validation.

**Table 1.** Data collections.

| Dataset | Role in study | Total N (patients) | ER+ (n) | ER- (n) | PR+ (n) | PR- (n) | HER2+ (n) | HER2- (n) |
| --- | --- | --- | --- | --- | --- | --- | --- | --- |
| TCGA-BRCA | Primary contrastive pre-training + downstream ER/PR/HER2 classification | 1098 | 807 | 291 | 714 | 384 | 192 | 906 |
| TCGA-OV | Supplementary dataset for contrastive pre-training | 585 | / | / | / | / | / | / |
| HistologyHSI-BC-Recurrence | Downstream test set | 47 | 38 | 9 | 32 | 15 | 10 | 37 |
| HER2 tumor ROIs - Yale HER2 cohort | Downstream test set | 192 | / | / | / | / | 93 | 99 |
| HER2 tumor ROIs - Yale trastuzumab response cohort | Downstream test set (auxiliary) | 85 | 66 | 18 | 62 | 22 | 85 | 0 |

Each sample may contain up to two WSIs: a diagnostic slide and a tissue slide. In total, 2,690 WSIs were included in this study, consisting of both 20x and 40x magnification images. Each WSI at the original resolution (level 0) was tiled into non-overlapping image patches of size W x H pixels, where W and H represent the patch width and height, respectively. In this study, both W and H were set to 256 pixels. During the tiling process, patches that did not contain tissue were discarded to reduce background noise. For each retained patch, the corresponding spatial coordinates (x, y) within the original WSI were recorded to preserve spatial information.

Bulk RNA-seq data were downloaded from TCGA and processed using standard normalization procedures. After normalization, housekeeping genes were removed to reduce the influence of constitutively expressed genes. The remaining genes were ranked according to their expression levels in descending order, and only the top 50 most highly expressed genes per sample were retained for downstream analysis.

### A frozen transcriptomic encoder guides contrastive pre-training of the slide-level vision encoder

GigaPath and BioBERT pretrained text encoder are aligned via a symmetric InfoNCE loss over patient-matched WSI-RNA pairs (**Fig. 1**). In the WSI branch, each H&E-stained WSI is tiled and encoded by the GigaPath tile encoder (ViT-Giant, pretrained via DINOv2 on 1.3 billion pathology tiles, kept frozen); the resulting patch embeddings are aggregated by the GigaPath slide encoder (dilated attention), which remains trainable throughout contrastive pre-training, to produce a slide-level representation *z*^*WSI*^ ∈ ℝ^*d*^. In the transcriptomic branch, bulk RNA-seq profiles are converted into a “gene sentence”: for each sample, the top 50 most highly expressed genes (after housekeeping-gene removal) are ranked by expression level and concatenated, in ranked order, into a single string of gene symbols, following gene-sentence formulation. This text sequence is tokenized and encoded by BioBERT pretrained text encoder, yielding a transcriptomic embedding *z*^*RNA*^; the text encoder is kept frozen throughout pre-training, serving as a fixed molecular reference against which the WSI encoder is aligned. Both *z*^*WSI*^ and *z*^*RNA*^ are projected by modality-specific heads *(g*^*WSI*^, *g*^*RNA*^) into a shared embedding space, where a symmetric InfoNCE loss with a learnable temperature (initialized at τ=0.07) pulls patient-matched pairs together while separating unpaired samples within each mini-batch. Following pre-training, the RNA encoder, both projection heads, and the contrastive loss are discarded; only the contrastively updated GigaPath backbone is retained. This backbone is evaluated on the downstream ER/PR/HER2 classification task under two protocols. In the first, a frozen-backbone linear probe, only a lightweight two-layer MLP classification head is trained on the fixed slide-level representation (our primary protocol). In the second, an end-to-end fine-tuned variant, the GigaPath backbone is further updated jointly with the classification head (secondary protocol). Together, these two protocols allow us to isolate the contribution of contrastive pre-training itself from that of downstream adaptation capacity.

**Fig 1.**
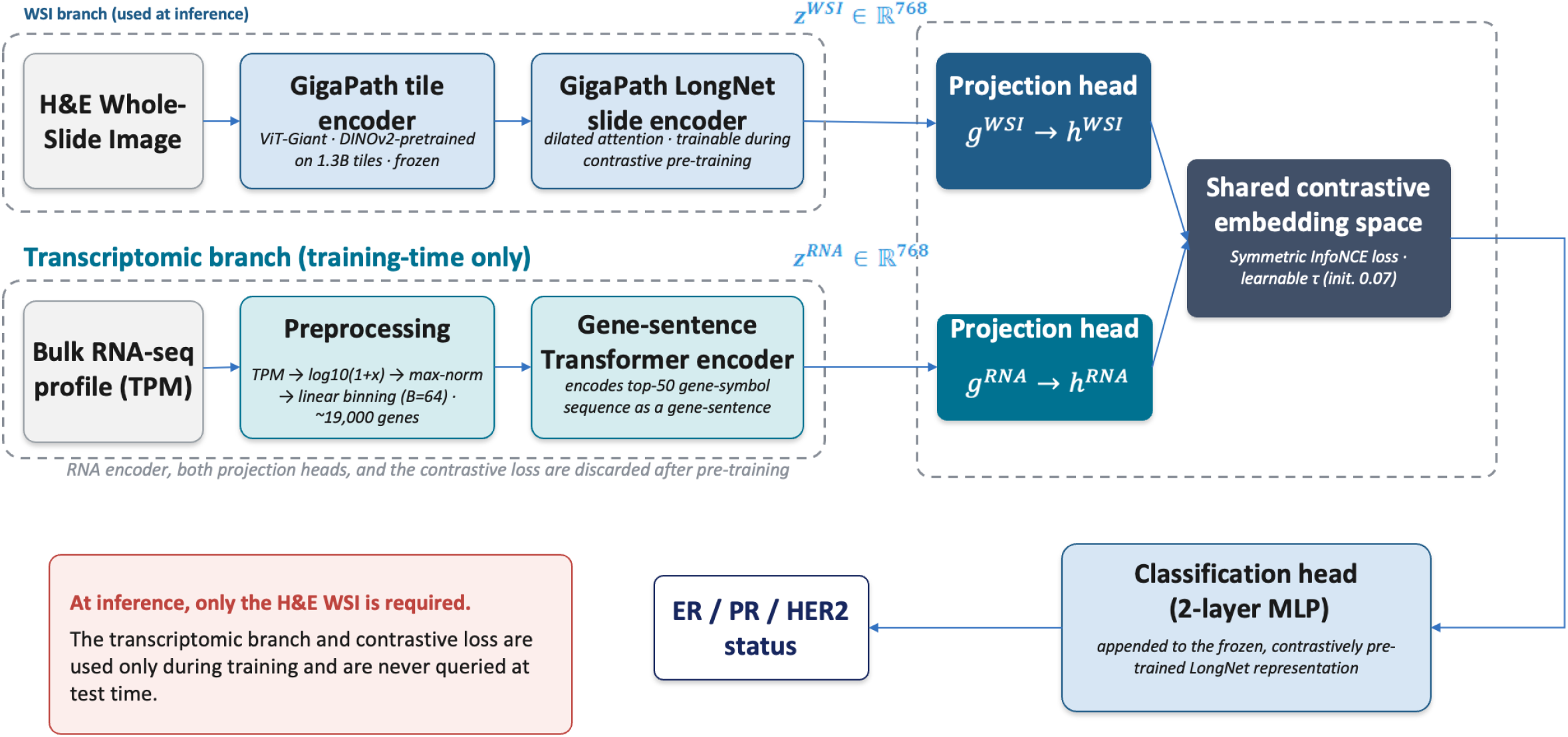
Schematic illustration of the overall design of the study.

### RNA-guided contrastive pre-training improves cancer subtyping

To test whether RNA-guided contrastive pre-training improves the WSI encoder’s ability to predict molecular subtypes, we compared two configurations, both trained and evaluated on TCGA-BRCA under the same 5-fold cross-validation splits: (i) the vanilla GigaPath baseline (no contrastive pre-training) and (ii) GigaPath contrastively pre-trained using a top-50-gene encoder. In this scheme, the top 50 most highly expressed genes per sample are ranked and encoded as a “gene sentence” text sequence, following the transcriptomic encoding protocol of BulkRNA-CLIP. Both configurations were evaluated under the frozen-backbone linear probe protocol, first via 5-fold cross-validation on TCGA-BRCA (**Fig. 2a-c**) and then on an independent, combined external test set (**Fig. 2d-f, Table 2**).

**Table 2.**
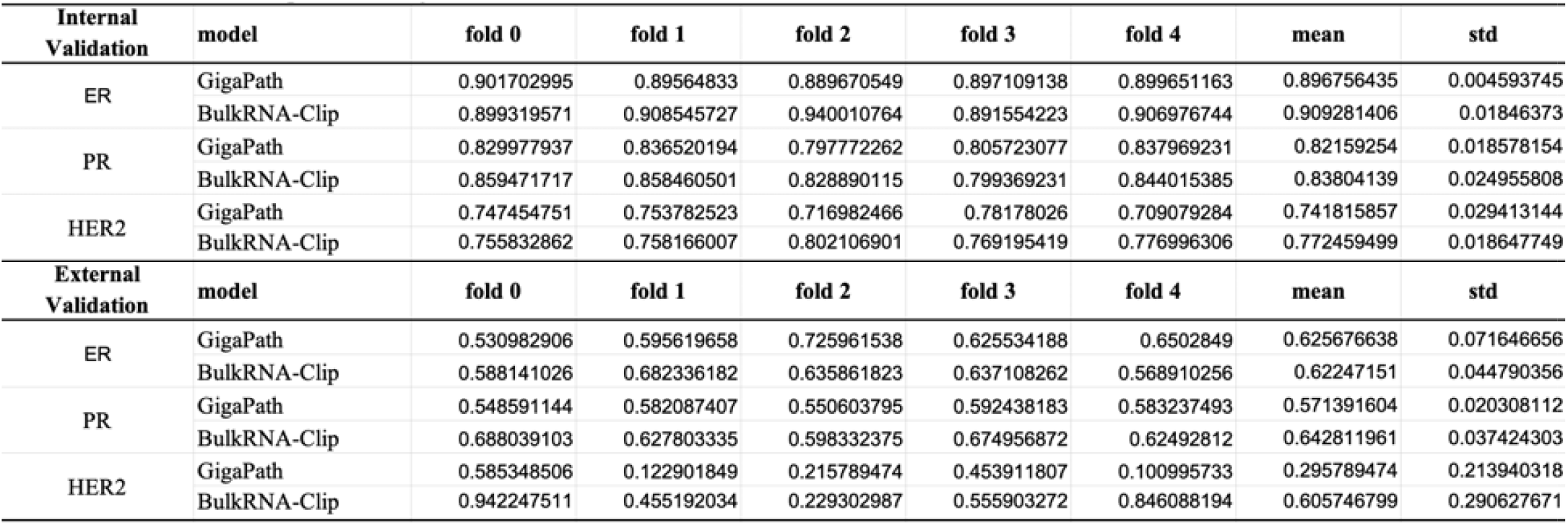
Performance comparison of GigaPath and BulkRNA-CLIP models for prediction of ER, PR, and HER2 status in internal and external validation cohorts.

**Fig 2.**
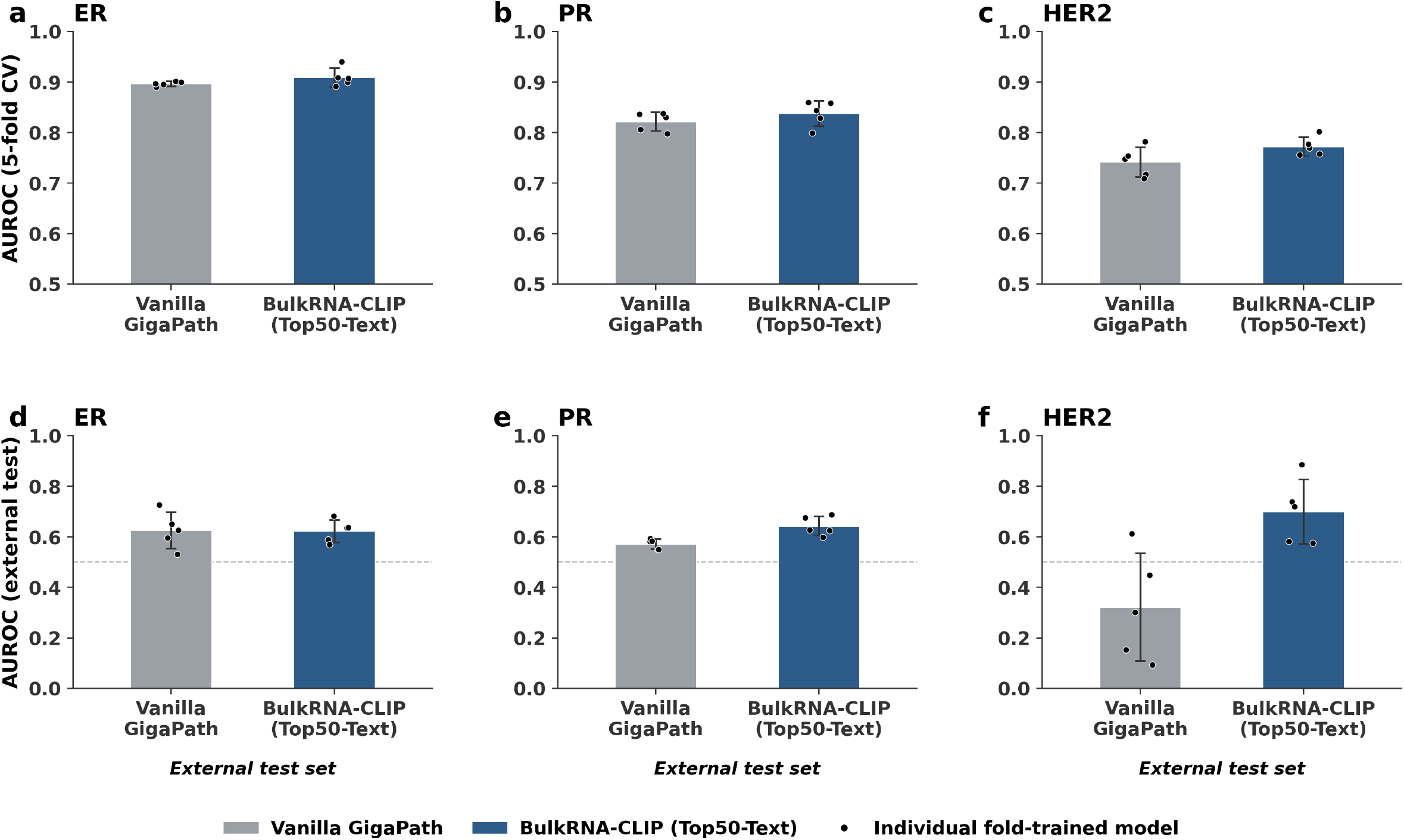
Performance comparison between Vanilla GigaPath and BulkRNA-CLIP models for prediction of ER, PR, and HER2 status. (a-c) AUROC scores from 5-fold cross-validation for prediction of (a) ER, (b) PR, and (c) HER2 status, comparing the Vanilla GigaPath model (gray) with the BulkRNA-CLIP model (blue). (d-f) AUROC scores on an independent external test set for prediction of (d) ER, (e) PR, and (f) HER2 status, comparing the same two models.

Under 5-fold cross-validation, BulkRNA-CLIP improved mean AUROC over the vanilla baseline for all three receptors: from 0.897 ± 0.005 to 0.909 ± 0.018 for ER (ΔAUROC = 0.013), from 0.822 ± 0.019 to 0.838 ± 0.025 for PR (ΔAUROC = 0.016), and from 0.742 ± 0.029 to 0.772 ± 0.019 for HER2 (ΔAUROC = 0.031) (**Fig. 2a-c, Table 2**). The direction of improvement was therefore consistent across all three receptors, with the largest gain observed for HER2. However, given the small number of folds (n = 5), the paired 95% confidence interval for ΔAUROC crossed zero for every receptor (ER: −0.015 to 0.040; PR: −0.004 to 0.037; HER2: −0.023 to 0.084), so these within-cohort gains are directionally consistent but not individually statistically significant.

On the combined external test set, the pattern was less uniform. AUROC was essentially unchanged for ER (vanilla baseline 0.626 vs. BulkRNA-CLIP 0.623, ΔAUROC = −0.003), improved for PR (0.571 to 0.643, ΔAUROC = 0.071), and improved substantially for HER2 (0.322 to 0.700, ΔAUROC = 0.379) (**Fig. 2d-f, Table 2**). The large apparent gain for HER2 was driven in part by high fold-to-fold variability in the vanilla baseline (SD = 0.213, including one fold with AUROC below 0.10), whereas BulkRNA-CLIP generalized more consistently across folds (SD = 0.129).

Together, these results indicate that RNA-guided contrastive pre-training with even a compact, top 50-gene transcriptomic representation can improve receptor-status classification from H&E images alone, without requiring any RNA-seq data at inference. Gains were consistent, if individually modest, under cross-validation on the training cohort, and more variable and receptor-dependent under external generalization, being most pronounced for HER2 and comparatively muted for ER (**Fig. 2**).

To visualize the threshold-dependent behavior underlying the AUROC values reported above, we additionally plot the fold-wise ROC curves for both configurations across all three receptors (**Fig. 3a-c**). Consistent with the AUROC improvements reported for 5-fold cross-validation, the ROC curves for BulkRNA-CLIP are shifted toward the upper-left of those for the vanilla GigaPath baseline across most of the false-positive-rate range for ER and PR, indicating a genuine improvement in discriminative ranking rather than one driven by a small number of folds or a narrow region of the curve. For HER2, the curves for BulkRNA-CLIP (Top50-Text) show tighter fold-to-fold clustering than the vanilla baseline, particularly at low false-positive rates, consistent with the higher overall AUROC reported in **Fig. 2c**. However, as the confusion-matrix analysis in **Fig. 4** shows, this improved ranking does not translate into higher sensitivity at the specific operating threshold examined, underscoring that AUROC and ROC-curve shape characterize ranking quality across all thresholds, not performance at the threshold actually used for classification.

**Fig 3.**
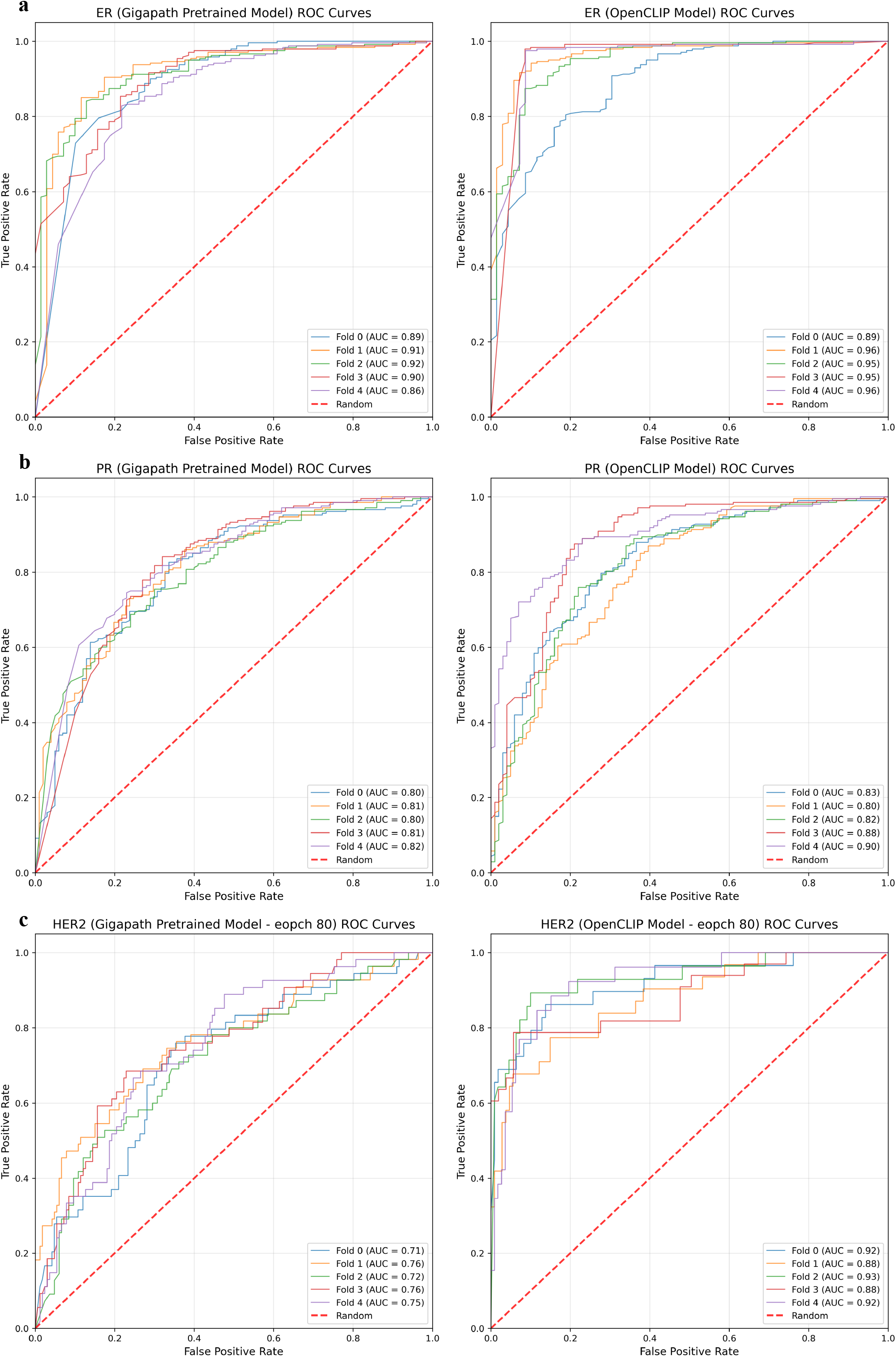
Fold-wise ROC curves for Vanilla GigaPath and BulkRNA-CLIP models. (a) ER, (b) PR, and (c) HER2 prediction tasks.

**Fig 4.**
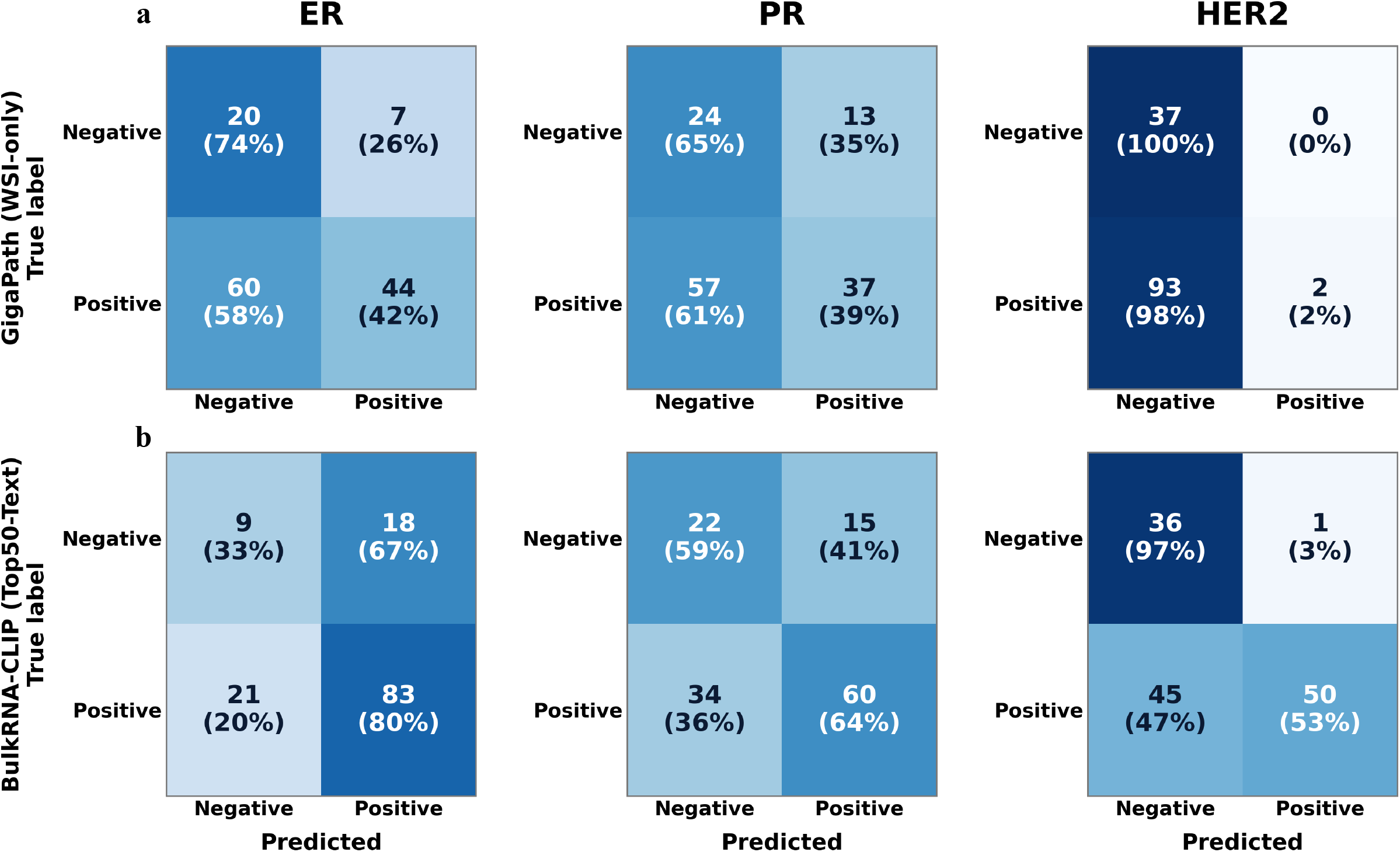
Confusion matrices comparing GigaPath and BulkRNA-CLIP models for prediction of ER, PR, and HER2 status. (a) Confusion matrices for the GigaPath model predicting ER (left), PR (middle), and HER2 (right) status. Rows indicate true labels and columns indicate predicted labels; each cell shows the number of cases and the corresponding row-wise percentage. (b) Confusion matrices for the BulkRNA-CLIP model predicting ER (left), PR (middle), and HER2 (right) status, presented in the same format as in (a).

### Contrastive pre-training modestly improves separation of receptor status in the learned slide representation

To ask whether contrastive pre-training reshapes the WSI encoder’s representation space in a way that better reflects molecular phenotype, beyond its effect on downstream classification accuracy, we computed the silhouette score of slide-level embeddings with respect to ER, PR, and HER2 status for the vanilla GigaPath baseline and for BulkRNA-CLIP at its best-performing contrastive pre-training checkpoint (epoch 19, the checkpoint used throughout Figs. 2-4). Under the vanilla GigaPath baseline, silhouette scores were negative for all three receptors (ER: −0.0011; PR: −0.0030; HER2: −0.0048 in **Fig. 5**), indicating that receptor-positive and receptor-negative cases are not separated by clinical label in the baseline embedding space; if anything, cases are marginally closer, on average, to the opposing label’s cluster than to their own. Contrastive pre-training with BulkRNA-CLIP shifted the silhouette score to positive values for all three receptors (ER: 0.0186; PR: 0.0123; HER2: 0.0003 in **Fig. 5**).

**Fig 5.**
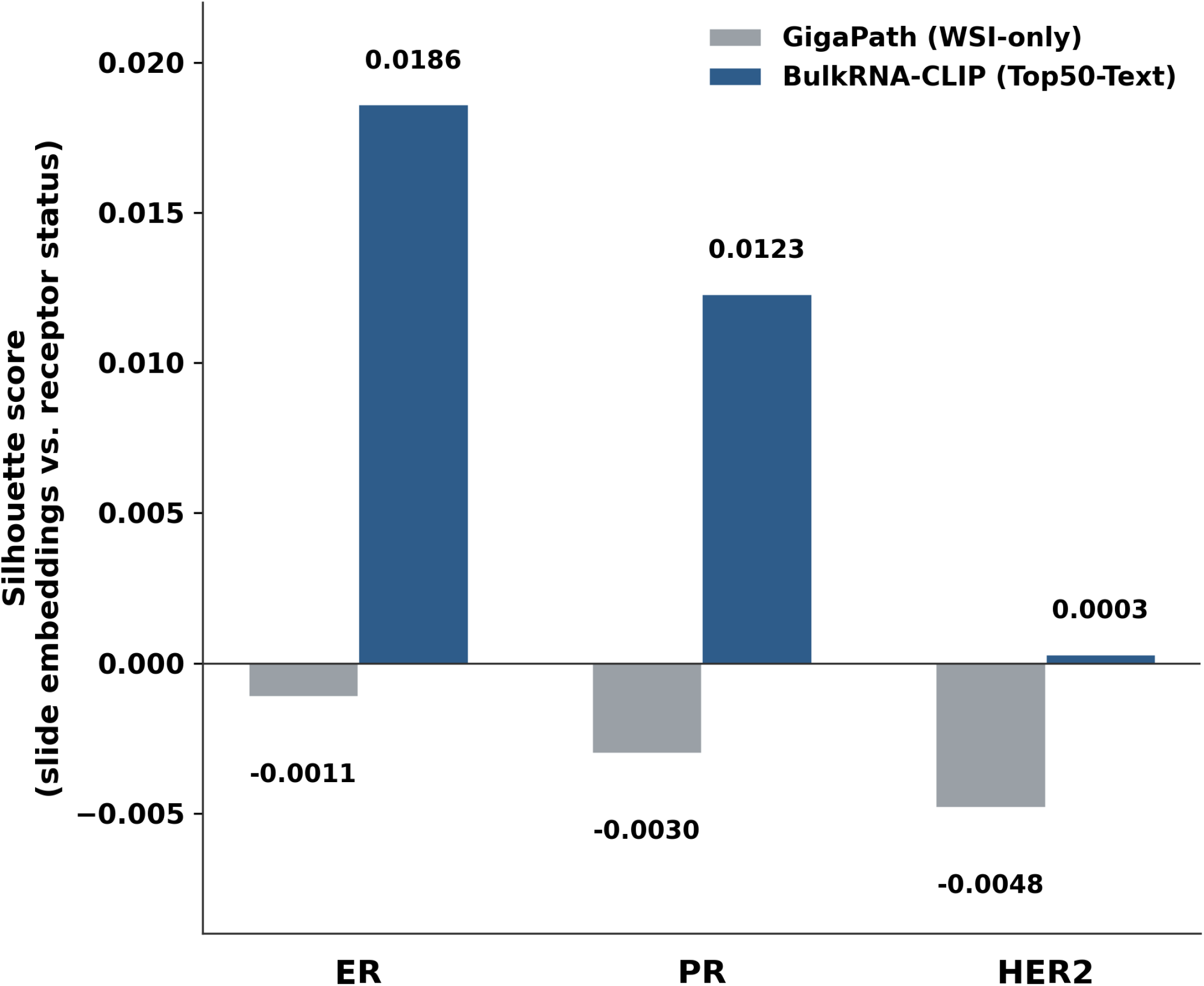
Silhouette scores of slide embeddings with respect to receptor status for GigaPath versus BulkRNA-CLIP models.

The magnitude of this shift should be interpreted cautiously: silhouette scores for both models remain close to zero in absolute terms (|score| < 0.02), far below the range (approaching 1) typically associated with well-separated clusters, indicating that receptor status is only weakly reflected in the global geometry of either embedding space. The consistent sign change from negative to positive across all three receptors nonetheless suggests that contrastive pre-training introduces a small but reproducible degree of receptor-relevant structure into the slide representation that is absent from the WSI-only baseline.

### RNA-guided contrastive pre-training shifts the sensitivity-specificity trade-off of receptor-status classification

Because AUROC summarizes classifier performance across all decision thresholds, it can obscure clinically relevant differences in the sensitivity–specificity trade-off at the operating threshold actually used for prediction. To examine this directly, we compared confusion matrices for the vanilla GigaPath baseline and BulkRNA-CLIP on an independent external cohort of 131 cases for ER and PR and 132 cases for HER2 (Fig. 4; a representative cross-validation fold is shown). All three receptors were imbalanced toward the receptor-positive class (ER: 104/131 positive, 79%; PR: 94/131 positive, 72%; HER2: 95/132 positive, 72%). For ER, the two models attained almost identical AUROC (0.626 vs. 0.622) yet occupied markedly different operating points: the vanilla GigaPath baseline recovered only 44/104 = 42% of positive cases at 20/27 = 74% specificity, whereas BulkRNA-CLIP recovered 83/104 = 80% of positive cases at 9/27 = 33% specificity. The same shift was seen for PR, where BulkRNA-CLIP raised sensitivity from 37/94 = 39% to 60/94 = 64% while specificity fell from 24/37 = 65% to 22/37 = 59%. Thus, for these two receptors RNA-guided contrastive pre-training moves the classifier toward the majority, receptor-positive class, trading specificity for sensitivity, a shift that is essentially invisible in the ER AUROC comparison.

For HER2, the gap between the two models was far larger and was not merely a threshold effect. The vanilla GigaPath baseline assigned nearly every case to the receptor-negative class, recovering only 2/95 = 2% of true HER2-positive cases at 37/37 = 100% specificity, and its mean AUROC (0.296) fell below chance. BulkRNA-CLIP recovered 50/95 = 53% of positive cases while retaining 36/37 = 97% specificity, with a mean AUROC of 0.606. The vanilla baseline’s degenerate, all-negative behavior combined with a below-chance AUROC indicates that its HER2 head carries essentially no transferable signal on this external cohort, rather than being merely mis-calibrated. These results reinforce the point that AUROC alone is insufficient: threshold selection and class-imbalance correction (e.g., threshold tuning per receptor, or reporting balanced accuracy and F1 alongside AUROC) warrant explicit attention before any of these models could be considered for clinical use, particularly for HER2, where per-fold AUROC varied widely (0.23-0.94 for BulkRNA-CLIP).

The residual difficulty of HER2 may reflect a specific limitation of the top-50-gene encoding strategy rather than a general failure of RNA-guided contrastive pre-training. Unlike ER and PR, whose transcriptional programs involve broad, multi-gene co-expression networks that are likely to be reflected among a sample’s most highly expressed genes, HER2-positive status is driven predominantly by focal ERBB2 gene amplification, a copy-number event rather than a broad transcriptional program. Because our top-50 gene selection ranks genes by absolute expression level rather than differential expression, a marker gene may fail to rank among the top 50 most highly expressed genes even in samples that are clinically positive for it, since the highest-expressed genes are typically dominated by keratins, ribosomal proteins, and other highly and ubiquitously expressed transcripts. Quantifying this directly in the TCGA training cohort, ERBB2 was present among the top-50 most highly expressed genes in 50.8% of HER2-positive patients, ESR1 in 29.5% of ER-positive patients, and PGR in only 4.6% of PR-positive patients, while all three appeared in ≤1% of the corresponding receptor-negative patients. The encoding is therefore highly specific but poorly sensitive: when a marker gene enters the caption it almost always indicates a true positive, but for most positive cases the contrastive learning objective receives no direct transcriptomic evidence of receptor status. ER retains an indirect route to supervision through co-expressed luminal genes that do enter the caption (SLC39A6, 53.5% of ER-positive vs. 2.9% of ER-negative patients; GATA3, 24.9% vs. 1.7%; STC2, 24.7% vs. 1.7%), whereas PR has by far the weakest direct signal of the three. This is a limitation that a full-transcriptome encoder such as BulkRNABert, which does not depend on absolute expression ranking, may be better positioned to address.

## Discussion

We show that a training-time-only, RNA-guided contrastive learning framework can improve receptor-status classification from H&E whole-slide images alone, using bulk RNA-seq profiles from TCGA-BRCA and without requiring any dedicated bulk-transcriptomics foundation model. Repurposing OmiCLIP’s gene-sentence encoding and pretrained text encoder, originally developed for spot-level spatial transcriptomics, for bulk RNA-seq profiles proved sufficient to yield a measurable, directionally consistent improvement over a vanilla, contrastive-pre-training-free GigaPath baseline across all three receptors (ER, PR, HER2) under 5-fold cross-validation (**Fig. 2a-c**), and a further, receptor-dependent improvement on an independent external test set (**Fig. 2d-f**). This supports the view, motivated in the Introduction, that the specific transcriptomic encoding strategy pioneered for spatial transcriptomics can be transferred to the far more widely available and substantially cheaper modality of bulk RNA-seq, without requiring a new encoder to be trained from scratch.

For classification tasks of this kind, we consider a consistent improvement in AUROC across independently trained models to be informative on its own, even when it does not reach conventional statistical significance. Genuine, clinically meaningful gains from a representation-learning intervention such as ours are typically on the order of a few AUROC points rather than tens of points; with only five cross-validation folds, effects of this magnitude are not expected to cross a 95% significance threshold under a paired test, so the absence of formal significance reflects the limited statistical power of a 5-fold design rather than evidence that the improvement is spurious. Under 5-fold cross-validation, BulkRNA-CLIP improved mean AUROC over the vanilla baseline for all three receptors, with the direction of improvement consistent across ER, PR, and HER2 and the largest gain observed for HER2 (**Fig. 2a-c**). We treat this consistency across three independently trained, receptor-specific classifiers, rather than the expected absence of significance at n = 5, as the more informative evidence that contrastive pre-training confers a real, if modest, benefit; a larger cohort or repeated cross-validation would be needed to establish significance formally. The silhouette-score analysis (**Fig. 3**) provides an independent line of evidence pointing in the same direction: although contrastive pre-training reproducibly shifts the geometry of the slide-embedding space toward better separation by receptor status, the absolute magnitude of this separation remains small (|score| < 0.02 for all receptors), implying that receptor status is only weakly and partially encoded in the resulting representation, consistent with a real but modest biological signal rather than a fully solved classification problem.

The improvement observed for ER and PR is consistent with the underlying gene biology captured by our top-50 highly-expressed-gene selection strategy. Estrogen receptor signaling drives a broad, multi-gene luminal differentiation program rather than a single focal event: ER-positive breast tumors characteristically co-express a coordinated set of luminal-lineage genes, including GATA3, FOXA1, XBP1, and luminal cytokeratins (e.g., KRT8, KRT18), alongside ER itself. Because these genes are both highly and consistently expressed in luminal-differentiated tumor cells, several of them are likely to be captured among a sample’s top 50 most highly expressed transcripts regardless of the exact expression-rank cutoff used, providing the contrastive objective with a redundant, multi-gene signal correlated with ER status rather than depending on any single marker gene. Progesterone receptor (PGR) is, moreover, itself a direct transcriptional target of estrogen receptor signaling, with its expression induced by ER pathway activity, so the same luminal transcriptional program that supports ER classification is directly mechanistically coupled to PR status, plausibly explaining why our top-50-gene encoding was similarly informative for both receptors. This broad, multi-gene, lineage-level signal stands in contrast to HER2, whose status is governed by a single focal copy-number event rather than a coordinated transcriptional program; despite achieving a higher AUROC than the vanilla baseline under cross-validation (**Fig. 2c**), BulkRNA-CLIP showed a severe loss of sensitivity for HER2 in our confusion-matrix analysis (**Fig. 5**), which we identify as a priority for follow-up work in the Future Directions discussed below.

Our study has additional limitations. First, all training and cross-validation were performed exclusively on TCGA-BRCA; while we evaluated generalization on several independent external test sets (Fig. 2d-f), the external test sets themselves vary in size, patient population, staining protocol, and scanning conditions relative to TCGA-BRCA, which may partly explain the divergent, receptor-dependent pattern of external-test performance we observed. Second, our current framework does not incorporate any transcriptomic data from other cancer types; TCGA ovarian serous cystadenocarcinoma is molecularly related to basal-like/triple-negative breast cancer^11,13^ and represents a natural candidate for a future cross-cancer contrastive pre-training extension of this work, which we leave for future study rather than reporting here. Third, our confusion-matrix and silhouette-score analyses were performed on a single held-out test partition and a single contrastive pre-training checkpoint, respectively, rather than being averaged across the full 5-fold cross-validation; extending these analyses across folds would provide a more robust estimate of their variability. Finally, several implementation details of our contrastive pre-training and downstream fine-tuning procedures (e.g., projection-head dimensionality, learning rate, batch size, and training duration; see Experimental Setup) remain to be finalized and reported.

Looking forward, we see several concrete directions building on the results presented here: (i) replacing the top-50, absolute-expression gene-sentence encoding with a full-transcriptome encoder such as BulkRNABert^14^ to test directly whether recovering ERBB2-level signal improves HER2 sensitivity without sacrificing the gains already observed for ER and PR; (ii) extending the cross-validation and external test-set evaluation to include balanced-accuracy, F1, and threshold-calibrated metrics alongside AUROC, particularly for HER2; (iii) revisiting the degree of adaptation applied to the GigaPath backbone during contrastive pre-training (e.g., parameter-efficient tuning versus full fine-tuning versus a frozen backbone), an axis we did not systematically vary in the results reported here; and (iv) extending this framework, once validated on TCGA-BRCA alone, to a cross-cancer contrastive pre-training design using TCGA ovarian cancer data, to test whether the molecular similarity between basal-like breast cancer and high-grade serous ovarian carcinoma can be exploited as an additional, complementary source of training-time transcriptomic supervision. Taken together, our results indicate that training-time-only, RNA-guided contrastive pre-training is a promising and practically deployable strategy for enriching WSI-based molecular subtype classification in breast cancer, while also identifying specific, mechanistically grounded limitations, particularly for HER2, that should be addressed before such a model could be considered for clinical use.

## Method

### Whole Slide Image Preprocessing

WSI preprocessing followed the standard pipeline established by GigaPath. Each whole-slide image was retrieved in digital format and subjected to automated tissue detection to exclude background, adipose tissue, and artifactual regions. Tissue-containing regions were segmented using Otsu’s thresholding applied to the luminance channel of the HSV color space. Qualified tissue tiles were then extracted at 40× magnification (0.25 μm/pixel) with a tile size of 256 × 256 pixels, with tiles containing less than 50% tissue coverage discarded. Each retained tile was independently encoded using the GigaPath tile encoder, a vision transformer (ViT-Giant) pretrained on over 1.3 billion pathology image tiles via a self-supervised DINOv2 objective^15^. This process yielded a sequence of 1,536-dimensional patch-level embeddings, the length of which varies across slides depending on tissue area. No stain normalization was applied, consistent with GigaPath’s training protocol, as the tile encoder has demonstrated robustness to staining variation across institutions. The resulting tile embedding sequences were subsequently passed to the GigaPath slide encoder to produce slide-level representations.

### Bulk RNA-seq Preprocessing

Bulk RNA sequencing preprocessing followed the “gene sentence” tokenization scheme proposed by OmiCLIP, applied here to bulk RNA-seq profiles from TCGA-BRCA rather than the spot-level spatial transcriptomics data used in OmiCLIP’s original framework. Raw RNA-seq counts were first normalized using transcript-per-million (TPM), followed by gene filtering to retain only protein-coding genes. To reduce dimensionality and mitigate the influence of ubiquitously expressed genes that carry limited tissue-specific information, we applied the following two-step selection strategy:

#### Housekeeping Gene Removal

Housekeeping genes, defined as genes with consistently stable expression across diverse tissues and conditions, were identified using the reference panel of the Housekeeping gene list used in OmiCLIP and excluded from further analysis. These genes, while essential for maintaining cellular homeostasis, provide minimal discriminative signal for tumor subtype classification and can introduce unwanted variance during cross-modal alignment.

#### Top-50 Gene Sentence Construction

Following housekeeping gene removal, the remaining genes in each sample were ranked by expression level in descending order, and the top 50 most highly expressed genes were retained and concatenated, in ranked order, into a single space-separated string of gene symbols (e.g., “ATP1A1 VTCN1 CTNND1 SLPI APP…”), following gene-sentence formulation. This text-based representation converts each sample’s transcriptomic profile into a short, natural-language-like sequence, capped at 76 tokens to match OmiCLIP’s original input length, that can be processed directly by a pretrained text encoder rather than requiring a dedicated numerical gene-expression encoder. The resulting gene-sentence string was tokenized and encoded by the transcriptomic (text) encoder described in Section X, initialized from BioBERT pretrained text-encoder weights, optionally with the text encoder locked during contrastive pre-training (see Contrastive Learning Objective), to produce the 256-dimensional transcriptomic embedding *z*^*RNA*^ used in the shared contrastive embedding space.

### Model Overview

H&E-stained WSIs provide rich morphological information, their standalone utility for predicting molecular subtypes of breast cancer, including ER, PR, and HER2 status, remains inherently limited. Integrating complementary modalities such as genomic or transcriptomic data represents a natural strategy to enrich the representation capacity of image-based models. However, multi-omics data including DNA methylation profiles and RNA sequencing are costly to acquire, technically demanding to process, and rarely co-available with histological slides in clinical or research cohorts. This asymmetry poses a fundamental bottleneck for direct multi-modal inference at deployment time. To address this challenge, we propose a training-time multi-modal contrastive learning framework that transfers transcriptomic knowledge into the WSI encoder without requiring RNA data at inference. The key insight is that bulk RNA-seq profiles, when available during training, can serve as privileged supervisory signals to shape a more biologically informed image representation, one that implicitly encodes molecular phenotype information otherwise invisible to morphology alone. At test time, only the H&E WSI is required, preserving clinical practicality. We refer to the resulting contrastively pre-trained WSI encoder as BulkRNA-CLIP throughout this paper, to distinguish it from the vanilla GigaPath baseline described under Comparison Baseline below.

### WSI Encoder

For whole-slide image encoding, we adopt GigaPath, a recently proposed foundation model for computational pathology that employs a hierarchical vision transformer architecture. Specifically, GigaPath utilizes a LongNet-based^16^ slide-level encoder capable of processing extremely long token sequences, making it well-suited to aggregate patch-level features across gigapixel WSIs. Each WSI is first tessellated into non-overlapping patches at 20× magnification, which are individually embedded by GigaPath’s tile encoder. The resulting patch token sequence is then passed through the LongNet slide encoder to produce a holistic slide-level representation *z*^*WSI*^ *∈ℝ*^*d*^.

### Transcriptomic Encoder

To encode bulk RNA-seq profiles, we adopt the transcriptomic (text) encoder from BioBERT. We implement transcriptomic branch represents each sample as a “gene sentence” (an ordered sequence of gene symbols) rather than operating on numerical gene expression values directly, an approach inspired by large language model-based single-cell methods such as GenePT^17^ and Cell2Sentence^18^. Following the preprocessing steps, the retained genes for each sample are ranked by expression level in descending order, and the top 50 most highly expressed genes are concatenated, in ranked order, into a single space-separated string of gene symbols (e.g., “ATP1A1 VTCN1 CTNND1 SLPI APP…”), capped at 76 tokens. Because this representation is a text sequence rather than a numerical vector, it is processed directly by a standard pretrained text encoder rather than requiring a dedicated gene-expression encoder architecture. This gene-sentence encoding scheme is borrowed from OmiCLIP. However, we do not use OmiCLIP’s own pretrained OpenCLIP-based text-encoder weights, since those weights were fine-tuned on Visium spot-level spatial transcriptomics data rather than on WSI-scale or bulk RNA-seq data. Instead, we initialize the transcriptomic encoder from BioBERT’s pretrained weights and use them directly, without further pre-training on bulk RNA-seq data, relying on the fact that the tokenization scheme (ranked gene symbols as text) is agnostic to whether the underlying profile derives from a Visium spot or a bulk RNA-seq sample. The encoder maps each gene-sentence token sequence to a sample-level transcriptomic embedding *z*^*RNA*^. In our work, the transcriptomic encoder is kept frozen throughout the cross-modal contrastive learning stage (see Contrastive Learning Objective below), serving as a fixed feature extractor of molecular phenotype information; we additionally evaluate a variant in which the text encoder is locked (frozen) versus left trainable during contrastive pre-training (see Experimental Setup). The resulting embedding *z*^*RNA*^ is projected into the same latent dimensionality as the WSI slide-level representation to facilitate cross-modal alignment.

### Contrastive Learning Objective

We frame the cross-modal alignment as a paired contrastive learning problem. For each training sample i, we have a matched pair 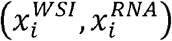, where the WSI and bulk RNA profile derive from the same patient. Slide-level and RNA-level representations are projected into a shared embedding space via modality-specific projection heads:

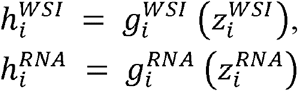

We optimize a symmetric InfoNCE loss [Oord et al., InfoNCE, 2018; Radford et al., CLIP, 2021] over a mini-batch of N paired samples. Intuitively, this objective encourages the WSI encoder to produce representations that are maximally aligned with the paired transcriptomic embedding while remaining discriminative against unpaired samples within the batch. The temperature parameter τ is initialized to 0.07 and updated during training.

During contrastive pre-training, both the GigaPath slide encoder and the projection head *g*^*WSI*^ .are updated end-to-end via gradient backpropagation. The RNA encoder weights are kept frozen throughout the contrastive pre-training stage, serving solely as a fixed feature extractor to provide molecular representations of transcriptomic inputs without participating in parameter optimization. This design choice is motivated by the fact that the text encoder has been extensively pretrained (first on LAION-5B as OpenCLIP, then fine-tuned by the authors on paired histology–transcriptomics data) and possesses strong gene-symbol semantic representation capability; freezing its weights effectively prevents overfitting on the limited paired breast cancer cohort while reducing computational overhead during training. While the transcriptomic encoder remains frozen, the WSI encoder is updated end-to-end via gradient backpropagation during contrastive pre-training, allowing GigaPath to learn morphological features predictive of the molecular phenotype captured by the frozen RNA embedding. Full end-to-end fine-tuning of a large pretrained vision backbone under limited paired supervision, however, carries a non-trivial risk of representation collapse or overfitting, a phenomenon also documented in the vision-language contrastive learning literature: Locked-image Tuning (LiT)^19^ found that keeping a strong pretrained image tower frozen during contrastive tuning can outperform allowing it to adapt jointly, particularly when paired training data is limited. Because GigaPath is itself a large-scale pretrained pathology foundation model rather than a randomly initialized network, we treat the degree of adaptation applied to it during contrastive pre-training as an explicit design axis rather than a fixed choice, and additionally evaluate parameter-efficient alternatives, namely IA3^20^ applied to GigaPath, and a fully frozen configuration in which only the projection heads *g*^*WSI*^ and *g*^*RNA*^ are trained, alongside full end-to-end fine-tuning, reporting whichever configuration yields the best-aligned representation on a held-out validation split.

### Downstream Task Fine-tuning

Following contrastive pre-training, the RNA encoder and projection heads are discarded entirely. Only the contrastively updated WSI encoder (GigaPath) is retained and fine-tuned for the downstream molecular subtype prediction tasks, namely ER, PR, and HER2 status classification. A lightweight task-specific classification head (a two-layer MLP with dropout) is appended to the slide-level representation. The entire pipeline, from patch embedding to classification, is optimized end-to-end using binary cross-entropy loss for each receptor status independently.

As our primary downstream protocol, the GigaPath backbone is kept frozen following contrastive pre-training, and only the lightweight classification head is trained on BRCA-train labels using binary cross-entropy loss for each receptor status independently. This linear-probing-style protocol ensures that downstream performance faithfully reflects the quality of the representation learned during contrastive pre-training, rather than being confounded by the additional adaptive capacity of a fully fine-tuned backbone. As a secondary, performance-oriented variant, we additionally report results with the LongNet backbone fine-tuned end-to-end alongside the classification head, to benchmark against prior work that does not hold the backbone frozen at this stage. This design ensures that the inference pipeline is strictly unimodal: at test time, only the H&E WSI is consumed, with no requirement for RNA-seq or any other ancillary molecular data.

### Comparison Baseline

To quantify the contribution of RNA-guided contrastive pre-training, we compare our approach against a vanilla GigaPath baseline: the publicly released GigaPath slide encoder, pretrained solely on WSI image data via a self-supervised objective (tile-level DINOv2 followed by slide-level masked autoencoding), without any exposure to RNA-seq or other molecular data, used directly for downstream molecular subtype prediction. The only difference between the two settings is therefore whether the WSI encoder additionally undergoes our RNA-guided contrastive alignment stage prior to downstream fine-tuning. The underlying GigaPath architecture and its WSI-only pretrained initialization are identical, and this baseline is evaluated under the same downstream protocol described above (a frozen-backbone linear probe as the primary comparison, and an end-to-end fine-tuned variant as a secondary comparison), with all other architectural components, training hyperparameters, and evaluation protocols held constant across settings. This controlled comparison isolates the added value of RNA-guided contrastive representation learning, on top of GigaPath’s existing WSI-only pre-training, for the WSI encoder’s predictive capacity in molecular subtyping.

### Experimental Setup

All models were implemented in PyTorch and trained on an NVIDIA A100 80GB GPU.

### Contrastive Pre-training

The GigaPath slide encoder was initialized from the publicly released GigaPath pretrained weights and fine-tuned during contrastive training. The transcriptomic encoder was initialized from the text encoder of BioBERT, which is a causal masking Transformer pretrained on LAION-5B. The transcriptomic encoder weights are kept frozen throughout the contrastive pre-training stage, serving solely as a fixed feature extractor to provide molecular representations of transcriptomic inputs without participating in parameter optimization. Both encoders were coupled with modality-specific two-layer MLP projection heads (hidden dimension: 768, output dimension: 768) that map representations into the shared contrastive embedding space, matching the embedding dimensionality used in text encoders. The InfoNCE loss was optimized with a learnable temperature parameter τ, initialized to 0.07. Training was performed using the AdamW optimizer with a learning rate of 1e-5, weight decay of 0.1, and a cosine annealing learning rate schedule with 500 warmup steps. The batch size was set to 64, and training proceeded for 20 epochs. Early stopping was applied based on contrastive loss on the validation set with a patience of 5 epochs.

### Downstream Fine-tuning

Following contrastive pre-training, the slide encoder was fine-tuned for each receptor prediction task (ER, PR, HER2) independently. A two-layer MLP classification head (hidden dimension: 256, dropout rate: 0.3) was appended to the slide-level representation. Training was conducted using binary cross-entropy loss, optimized with AdamW (learning rate: 2e-3, weight decay: 0.01) for 20 epochs with early stopping (patience: 10 epochs) monitored on validation AUROC. To address class imbalance, positive class weighting and oversampling were applied. The same fine-tuning protocol was applied identically to the baseline GigaPath model (without contrastive pre-training) to ensure a fair and controlled comparison.

